# Genetics and epigenetic alterations of hexaploid early generation derived from hybrid between *Brassica napus* and *B. oleracea*

**DOI:** 10.1101/148940

**Authors:** Qinfei Li, Zhiyong Xiong, Jiaqin Mei, Hongyuan Song, Wei Qian

## Abstract

Good fertility was observed previously in hexaploid derived from hybrid (ACC) between calona ‘Zhongshuang 9’(*Brassica napus*, 2n = 38, AACC) and kale ‘SWU01’ (*B. oleracea var. acephala*, 2n = 18, CC). However, the mechanism to underlying the character is unknown. In the present study, genetic and epigenetic alterations of S0, 6 S1, and 18 of their S2 progenies with hexaploid chromosome conformation (20A + 36C) were selected to compare with ACC and their parental species. 13.08% and 26.45% polymorphism alleles different from two parental species were identified in ACC via 58 SSR (simple sequence repeats) and 14 MSAP (methylation sensitive amplified polymorphism), respectively. 33.74% new alleles in DNA methylation, but not in DNA sequence were detected in S0 after chromosome doubling of ACC. DNA profilling revealed a little genetic but much epigenetic differences among S0, S1 and S2 generations. Genetic alteration was relatively stable, because only 8.09% and 3.21% alleles inheriated from ACC were changed in S2 and S1, respectively. While on average of 52.44 ± 5.32% DNA methylation site inherited from ACC were detected in S1, and 41.52 ± 9.04% in S2 due to dramatic epigenetic variance among early generations. New DNA methylation sites occurred in S0 would inheritated into the successive generations, but the frequency was decreased because some new site might be recovered. It demonstrated that much DNA methylation but a little DNA sequence variance was occurred in hexaploid early generation.

## Introduction

*Brassia napus* (2n = 38, AACC) is one of the most important oilseed crop in the world. It was derived from interspecfic hybridization between *B. rapa* and *B. oleracea* with natural chromosome doubling (U 1935). Its genetic basis was narrower than parental species due to its short history of domestication and intensive mordern breeding (Becker et al. 1995; Girke et al. 2012; Seyis et al. 2003). Introgressing genetic components of parentel species plays an important role in broadening its genetic basis. Previously, new strategy of using hexaploid (A^n^A^n^C^n^C^n^C^o^C^o^) derived from *B. napus* (A^n^A^n^C^n^C^n^) and *B. oleracea* (C^n^C^n^) was developed, and as bridge to cross with *B. rapa* to produce new type *B. napus* was created, and it showed new type *B. napus* owned high potential to broaden genetic basis of natural *B. napus* (Li et al. 2013). In case promoting this strategy in *B. napus* breeding program, hexaploid stability was the key factor. Previously, high frequency of euploid and good fertility was found in the progenies of the hexaploid, but the mechanism to underly the phenomoneon was unknown.

In nature, polyploidy is common, and it plays an important role in the evolution and domestication of several major crops, such as wheat, cotton and rapeseed (Chen 2010; Olsen and Wendel 2013; Chalhoub et al. 2014). During the process of polyploidy, genetic and epigenetic alterations, referred as ‘genomic shock’ often occurred (McClintock 1983), resulting in diverse genetic structure and gene function. In resynthesized polyploid, the genetic structure alterations including gene sequence addition and DNA sequence diversity, may result in chromosome recombination, chromosome rearrangement and rapid DNA sequence elimination (Akhunov et al. 2013; Gaeta et al. 2007; Xiong et al. 2011). And epigenetic changes, such as DNA methylation, histone modification and retrotransposon activation, might result in gene silencing or activation (Kantama et al. 2013; Sarilar et al. 2013). Hexaploid (A^n^A^n^C^n^C^n^C^o^C^o^) was derived from hybrid between *B. napus* and *B. oleracea*, it might induce further genomic variations involving change in DNA sequences and epigenetic modifications in the progenies. To verify this, we chose 6 S1 and their 18 S2 lines with hexaploid chromosome conformation to characterize genetic and epigenetic alteration via simple sequence repeats (SSR) and methylation-sensitive amplification polymorphism (MSAP) analysis, and found more DNA methylation alterations than SSR variance in the early generation of hexaploid.

## Results

### Development of hexaploid S1 and S2

In previous study (Li et al. 2013), the hexaploid S0 derived from hybrid between *B. napus* (‘Zhongshuang 9’) and *B. oleracea* var. *acephala* (‘SWU01’) exhibited good fertilization and high frequency of euploid in pollen mother cells, its bigger size flower could be used as indicator of hexaploid individuals in the progenies (Fig. 1A-D). Here we choose 21 individuals of S1 with big size flower to check their chromosome conformation, and found all of them had karyotype of AACCCC identified by FISH. Subsequently, 77 S2 were randomly chosen to check chromosome conformation, and 56 individuals were identified with hexaploid (Fig. 1E-F). Those findings were in accordance with previous observation (Li et al. 2013).

**Fig. 1.**
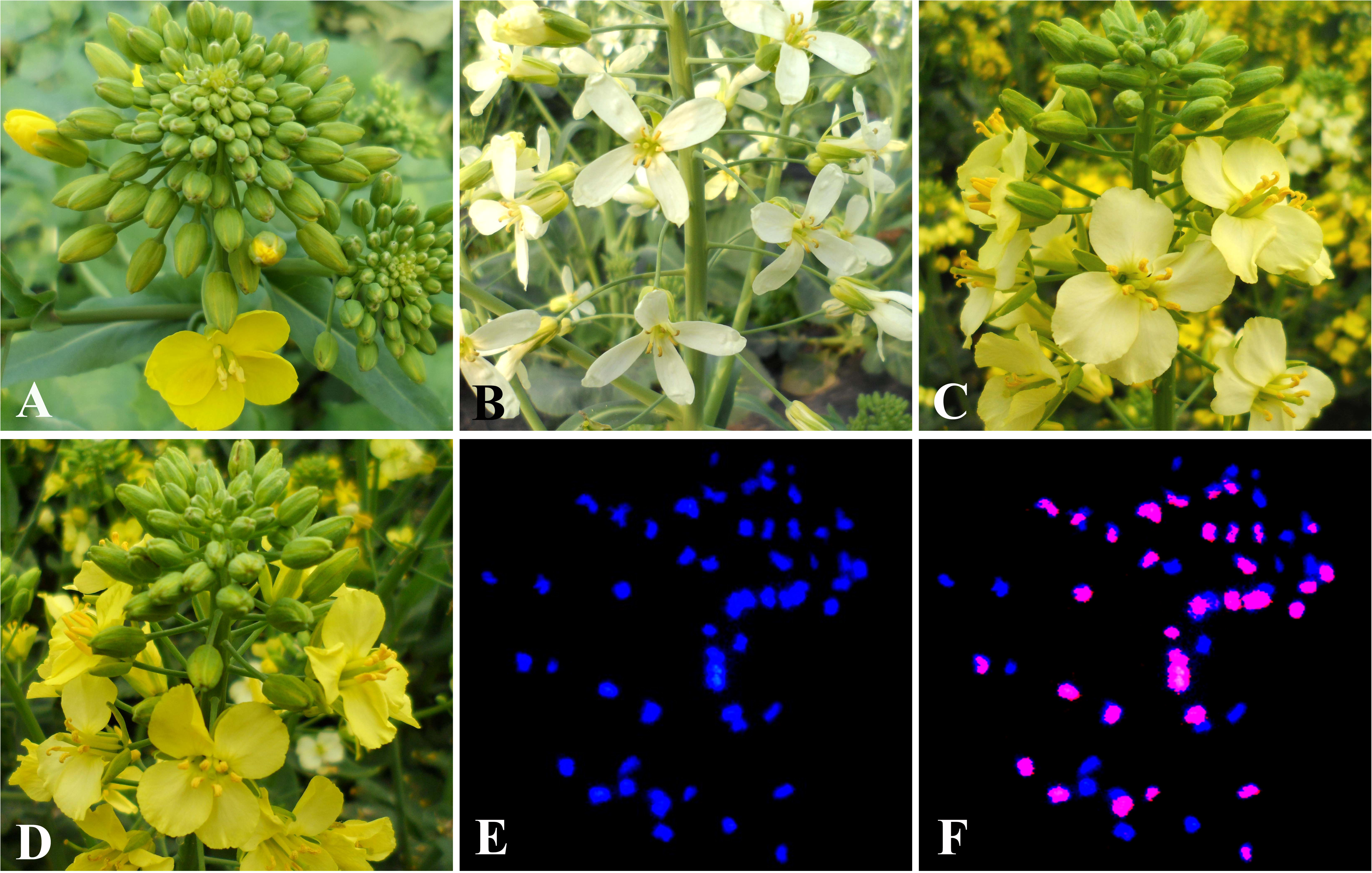
Morphology and chromosome conformation of hexaploid. **A** Flowers of *B. napus* (Zhongshuang 9); **B** Flowers of *B. oleracea* (SWU01); **C** Flowers of hexaploid S0; **D** Flowers of one individual hexaploid S2; **E-F** One tissue cell with chromosome conformation (36C + 20A), in which, chromosomes were counterstained with 0.2% 4’-6-Diamidino-2-phenylindole solution (blue) and the C chromosomes were identified with BoB014O06 fluorescence in situ hybridization signals (red).

The individuals owing the hexaploid conformation were called as euploid, while individuals lossing or gaining chromosmes were callled as aneuploid. In the present study, it is found that the average seed set was relatively stable in the early successvie generations of hexaploid (S0, S1 and S2). The average seed set of open pollination was significantly more than self-pollination in both S1 and S2 generations (*P* < 0.05), but no significant differences were found for average seed set between S1 and S2 generations. The mean value of aneuploid was a little more than euploid by calculating seed set of 27 S1 (21 euploid and 6 aneuploid) and 78 S2 (57 euploid and 21 aneuploid) individuals (Fig. 2).

**Fig. 2.**
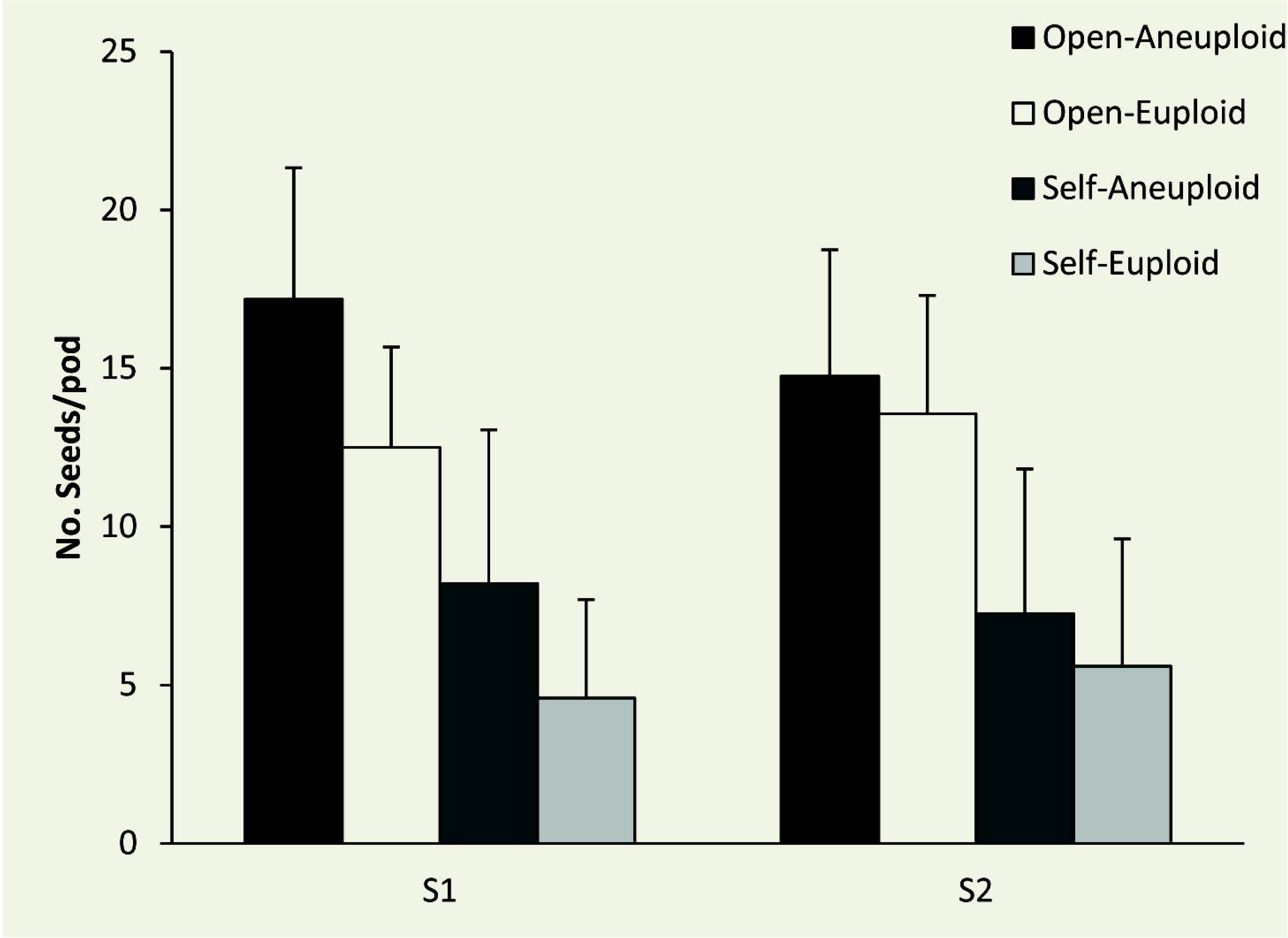
Seed set of hexaploid 27 S1 (21 euploid and 6 aneuploid) and 78 S2 (57 euploid and 21 aneuploid).

### Interspecific hybridization and chromosome doubling leading to genetic and epigenetic variance in AACCCC S0

To clarify genetic and epigenetic variance in the early generation of hexaploid,18 S2, 6 S1 with hexaploid chromosome conformation were compared with triploid ACC, S0 and parental species via SSR and MSAP, respectively. In total, 130 polymorphic alleles were amplified by 58 SSR markers. Comparing with parental species, triploid ACC shared 33 (25.39%) alleles with two parents, 80 (61.53%) alleles to a single parent, but had 17 (13.08%) unique alleles. No variance was detected between S0 and ACC via SSR.

Interspecific hybridization brought less genetic variance than epigenetic variance in ACC comparing to two parents, and chromosome doubling triggered epigenetic but no genetic changes in S0 in comparison with ACC. A total of 246 polymorphic loci were detected by fourteen primer combinations via MSAP. The variance of cytosine methylation was 26.45% between ACC and its two parents, while the variance between S0 and ACC was 33.74%. It indicated that the variation of DNA methylation in AACCCC S0 was caused by interspecific hybridization between *B. napus* and *B. oleracea* and chromosome doubling in the tissue culture.

### DNA sequence and DNA methylation variance of hexaploid S1 and S2

It was found that variance of DNA sequence and cytosine methylation ocurred in the early hexaploid generations S1 and S2 clarified by SSR and MSAP. It was found that S2 showed more variance than S1 in DNA sequence and DNA methylation alteration, and the average DNA methylation variance was significantly more than that of DNA sequence in each generations (*P* < 0.01). The average DNA sequence variance between S2 and S1 was 8.48 ± 4.94%, little higher than that between S1 and S0 (3.00 ± 1.45%). While the average variance of 5’CCGG site methylation between S2 and S1 generation (40.65 ± 7.14%) was more than that between S1 and S0 (34.62 ± 4.80%) (Table 1).

**Table 1.**
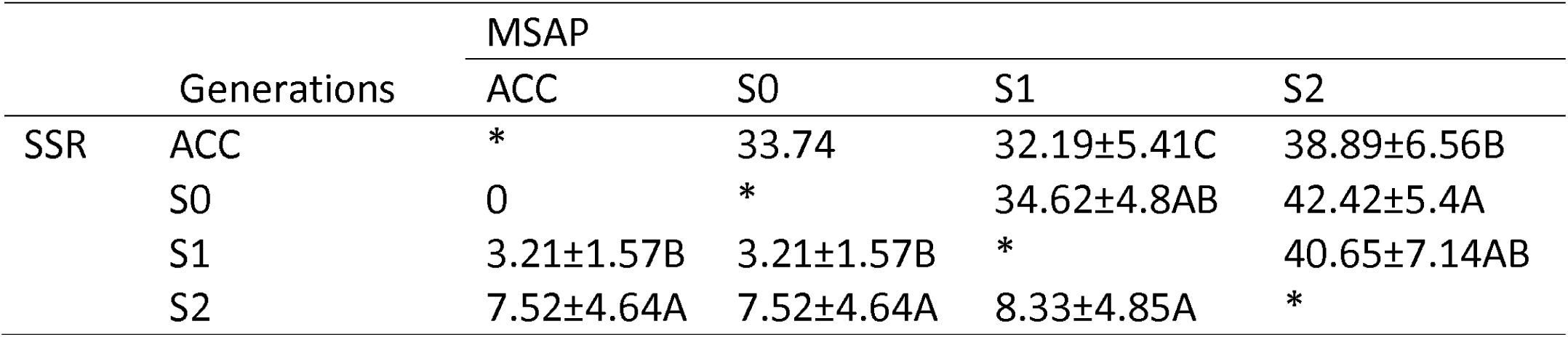
Genetic and epigenetic variance among ACC, S0, S1 and S2 detected by SSR and MSAP.

Herein, 9 S2 and 4 S1 were clustered together with S0, ACC and *B. napus*, while 7 S2 were close to *B. oleracea* in the phylogenetic tree constructed by polymorphic loci of MSAP (Fig 3). In the S1 lines, the DNA methylation variance of I2 (26.45%) was the lowest, while I4 (39.43%) was the most than others, and it showed highest variance to its offspring (47.83 ± 3.41%) (Table 2). It demonstrated that the variance of DNA methylation occurred more dramatically than DNA sequence in the early successive generation of hexaploid.

**Fig. 3.**
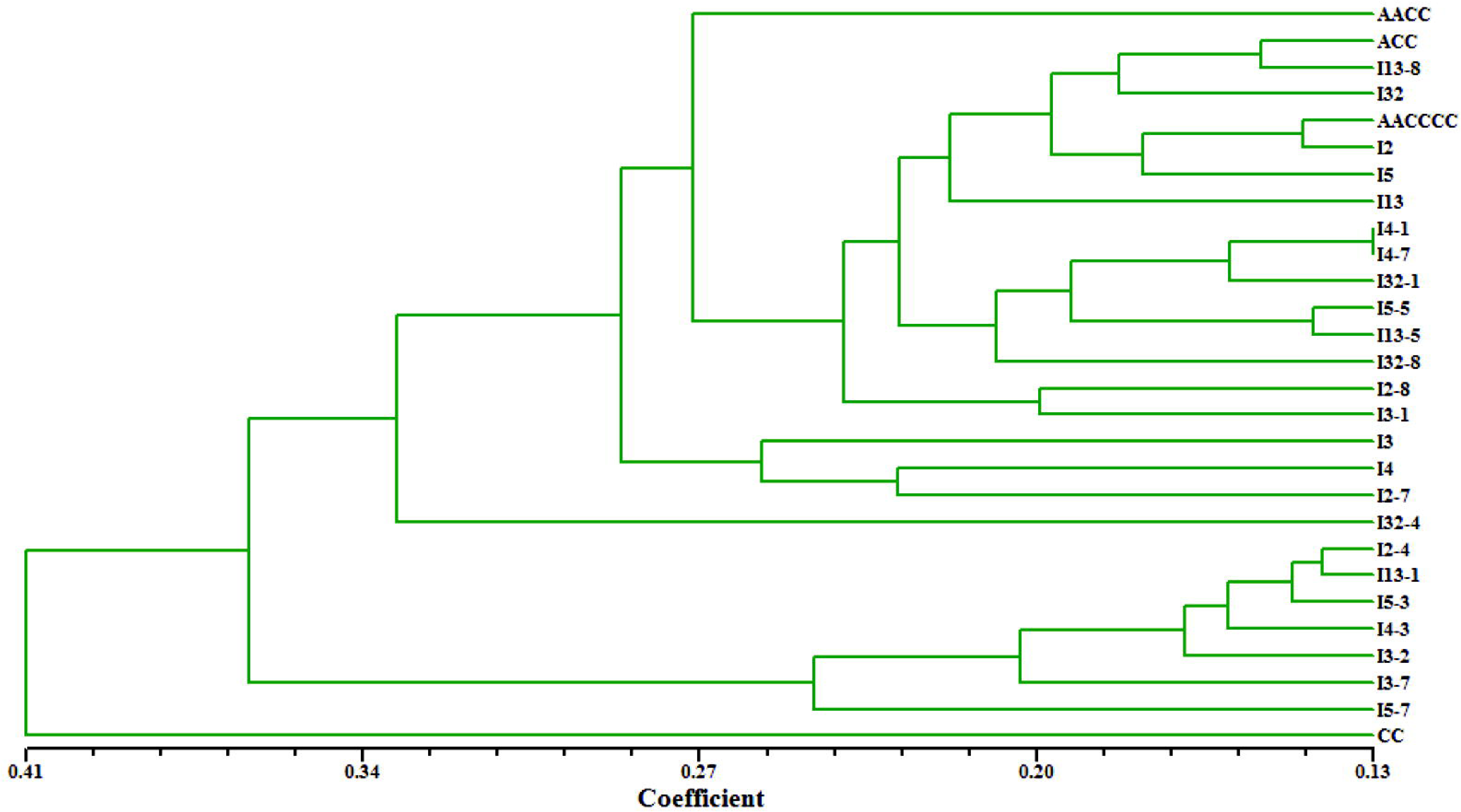
Cluster of hexaploid S0, S1 and S2, comparing with *B. napus* (AACC) and *B. oleracea* (CC) by MSAP

**Table 2.**
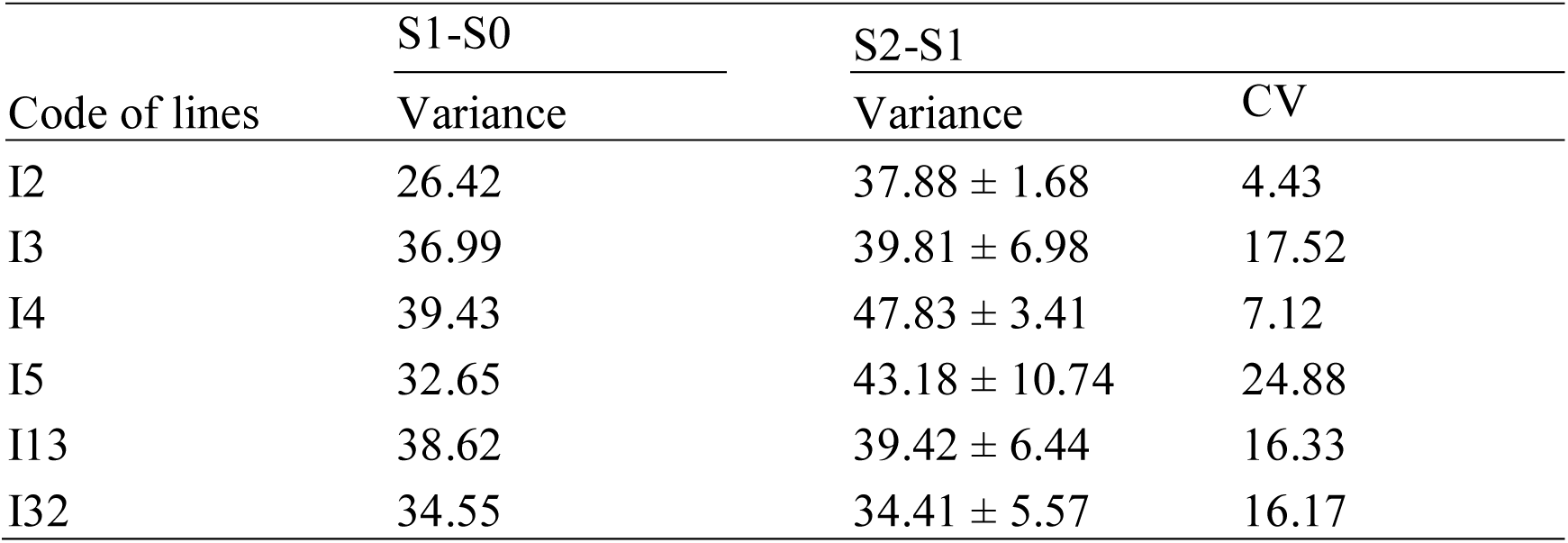
DNA methylation variance among S0, S1 and S2 generation

| Code of lines | S1-S0 | S2-S1 | CV |
| --- | --- | --- | --- |
|  | Variance | Variance |  |
| I2 | 26.42 | 37.88 ± 1.68 | 4.43 |
| I3 | 36.99 | 39.81 ± 6.98 | 17.52 |
| I4 | 39.43 | 47.83 ± 3.41 | 7.12 |
| I5 | 32.65 | 43.18 ± 10.74 | 24.88 |
| I13 | 38.62 | 39.42 ± 6.44 | 16.33 |
| I32 | 34.55 | 34.41 ± 5.57 | 16.17 |

### Genetic and epigenetic inheritation

The genetic and epigentic alteration of ACC could inheritated into S0 and its offspring, but less epigenetic alteration were inheritated due to its dramatic variance. In the early generation of hexaploid AACCCC, all the genetic components from ACC were inherited into AACCCC S0, on average of 97.00 ± 1.45% genetic components were inherited into S1 generation, then 89.86 ± 5.23% were inherited into the successive S2 generation according to the DNA sequence variance via SSR. However, it was found that only 66.26% 5’CCGG site methylation originated from ACC were inheritated into S0, on average of 52.44 ± 5.32% were inherited into S1, and 41.52 ± 9.04% were inherited into S2 successively.

Comparing to the variance calculated by all the DNA methylation sites among hexaploid S1 and S2 generations, the variance of inherited DNA methylation sites was less, and some new DNA methylation sites in S1 might be recovered in the S2 generation (Fig. 4). For the new 5’CCGG site methylation of S0, on average of 36.14 ± 4.94% was detected in S1, 16.10 ± 3.35% were detected in S2; while for the new DNA methylation sites in S1 differenting from S0, on average of 41.89 ± 9.36% was detected in S2. It showed that DNA methylation was inheritated into the successive generations, new DNA methylation occurred in S0 would inherited into successive generations, but the inherited DNA methylation were decreased while variance were increased in the next generation.

**Fig. 4.**
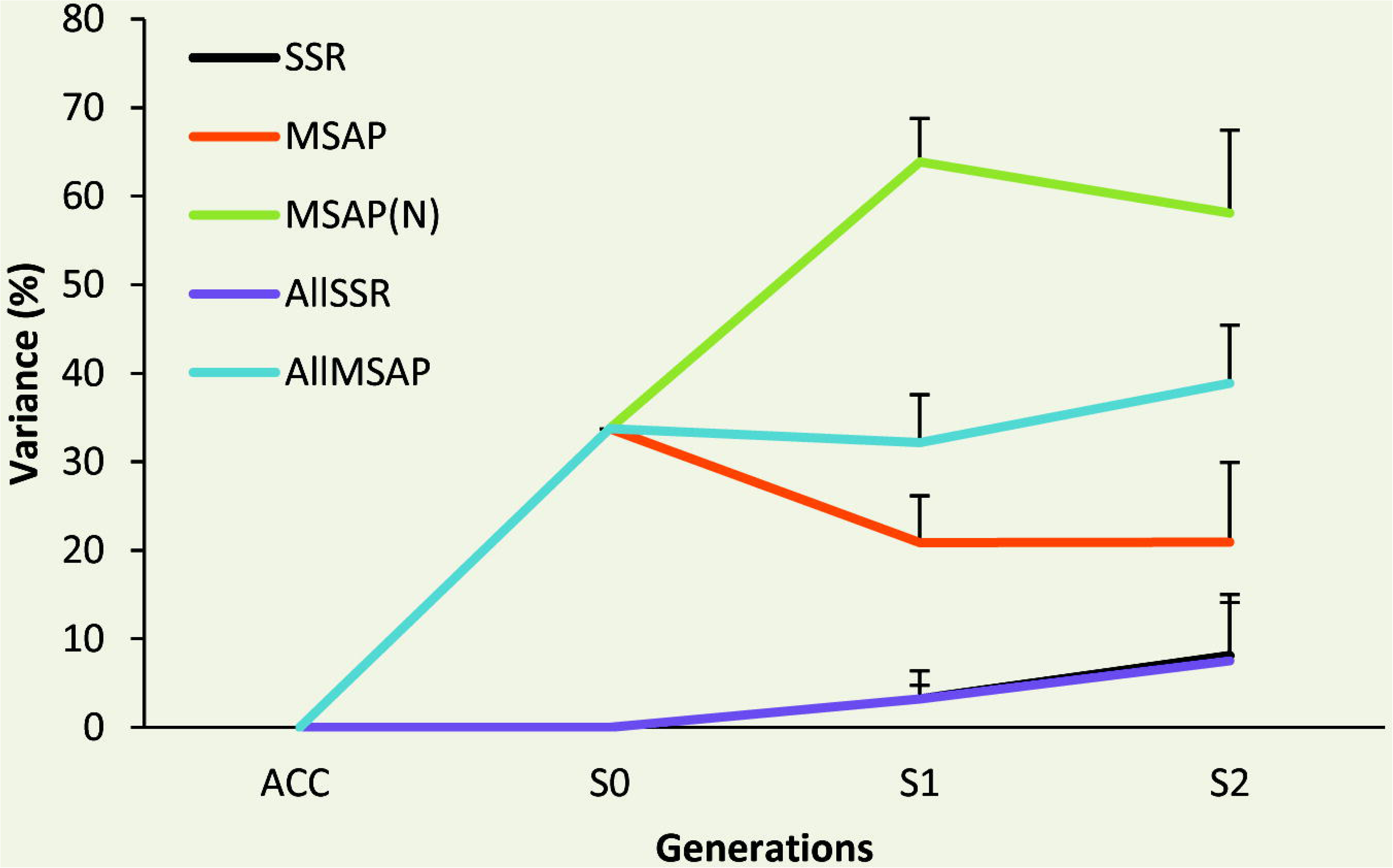
Genetic and epigenetic variance among ACC, S0, S1 and S2 detected by SSR and MSAP. AllMSAP and AllSSR was the variance of all DNA methylation sites and DNA among generations, respectively; MSAP and SSR was the variance of inheritated DNA methylation and DNA sites from ACC; MSAP (N) was the variance of new pattern of DNA methylation occurred in S0.

## Discussion

In this study, DNA sequence and DNA methylation were changed in the ACC hybrid comparing to parental species, and new pattern of DNA methylation but no DNA variance were occurred in hexaploid S0 (AACCCC) after chromosome doubling, which might caused by interspecific hybridization and polyploidization. During the process of allopolyploidy, ‘genomic shock’ of genetic and epigenetic variance occurred due to its genome experiences extensive and rapid genome restructuring (Akhunov et al. 2013; Gaeta et al. 2007; Soltis and Soltis, 2009; Wolfe 2001; Xu et al. 2009). For the plant genome evolution, the major genetic variance was lossing duplicated DNA sequence from allopolyploid parental species (Langham et al. 2004), while cytosine methylation is a stable mark in epigenetic variation and is important for diverse epigenetic phenomena (Bird 2002).

It has been reported that interspecific hybridization and allopolyploidization can induce more changes in DNA methylation than genetic changes in diverse plant taxa (Hegarty et al. 2011). For example, in naturally formed *Spartina* allotetraploids there were little genetic changes; while changes in DNA methylation occurred in > 30% loci assessed (Salmon et al. 2005). In euploid individuals of several synthetic allohexaploid wheats, rapid genetic changes were rare or nonexistent, but DNA methylation repatterning occurred extensively (Zhao et al. 2011). Even in natural populations, vary more in DNA methylation than in DNA sequence were shown by MSAP analyses (Lira-Medeiros et al. 2010; Salmon et al. 2008). In accordance with previous study, changes in DNA methylation also was more than that of DNA sequence in the early generation of hexaploid derived from interspecific hybridization between *B. napus* and *B. oleracea*, that is, their epigenetic diversity is higher than their genetic diversity.

It is well established that cytosine methylation, is often heritable to organismal progenies and the DNA methylation sites might undergo frequent reversions (Boyko and Kovalchuk 2011; Sani et al. 2013). Previously study showed that new pattern of cytosine methylation was commonly occurred in new allopolyploids comparing to parental species (Lukens et al. 2006; Xu et al. 2009; Zhao et al. 2011), and the changes were further exaggerated after genome doubling (Xu et al. 2014). In the early generation of resynthesized *B. napus*, high DNA methylation and little genomic variation were detected, and the DNA methylation was inherited into progenies (Gaeta et al. 2007; Lukens et al. 2006). In the isogenic *Arabidopsis* plants, epialleles occurred at a much higher frequency than genetic mutations, and the DNA methylation sites undergo frequent reversions by wholegenome BS-Seq (Becker et al. 2011; Schmitz et al. 2011). In the present study, cytosine methylation changes were occurred in ACC hybrid due to interspecific hybridization between *B. napus* and *B. oleracea*, and these variation was inheritated into S0, S1 and S2 via chromosome doubling and successive self-pollination.

Polyploidy is a driving force in genome evolution and plant domestication. In natural, in order to adapt different environment, plants often show striking phenotypic adaptations. Although phenotypic diversity in plants was thought to be attributable to genetic variation, epigenetic variation may also contribute to plant adaptation and other evolutionary processes. In the resynthesized allopolyploid *Spartina anglica*, phenotypic variability could be explained by epigenetic variability, rather than genetic variability (Baumel et al. 2001; Parisod et al. 2009). And it also hypothesized that epigenetic variation having an impact on the ecology and evolution of natural populations, if only epialleles arised in populations must be inherited with sufficient stability (Cubas et al. 1999; Manning et al. 2006). In the present study, although not significant correlation between fertility and genetic variance or epigenetic variance was detected, relative stable genetic variance was occurred among generations, and it might contributed to the relative stable fertility. High frequency of DNA methylation changes was detected among generations, but part of DNA methylation patterns were inherited. We hypothesize that stable epigenetic alleles might be included in these inherited DNA methylation alleles, and it might be correlated with the stable fertility,eand they might contributed to the stability of hexaploid due to the DNA methylation alleles inheritation.

## Materials and methods

### Plant materials

The hexaploid (A^n^A^n^C^n^C^n^C^o^C^o^) was developed by chromosome doubling of triploid hybrid (ACC) derived from hybrid between *B. napus* (‘Zhongshuang 9’, 2n = 38, AACC) and *B. oleracea* var. *acephala* (‘SWU01’, 2n = 18, CC) in previous study (Li et al. 2013). It was successively self-pollinated to develop S1 and S2 generations. Seed set and chromosome number of S1 and S2 lines were checked. And 6 S1 and 18 S2 with hexaploid chromosome conformation were chose to detect genetic and epigenetic alteration via simple sequence repeat (SSR) and methylation-sensitive amplification polymorphism (MSAP), respectivly.

### Chromosome conformation identification

To investigate the chromosome conformation of the hexaploid, fluorescent in situ hybridization (FISH) was performed according to the protocol of Leitch and Heslop-Harrison (1994) with some modifications. The probe was developed by extracting DNA of the C-genome-specific BAC clone, BoB014O06, which can hybridize with the C chromosomes, but not with the A chromosomes in *Brassica* (Howell et al. 2002), and labeling with Bio-11-dUTP by random priming. The hybridization signals of the BAC clone probe were detected by Cy3-labeled streptavidin (Sigma, USA), and chromosomes were counterstained with 0.2 % 4’-6-Diamidino-2-phenylindole (DAPI) solution (Roche, BaseI, Switzerland), mounted in antifade solution (Vector) and examined under a fluorescent microscope (Nikon Eclipse 80i, Japan) equipped with CCD camera. Images were processed using the software of Adobe Photoshop version 8.0.

### SSR and MSAP fingerprinting

The genomic DNA of S0, S1 and S2 with the chromosome conformation of hexaploid, and parental lines were isolated from young leaves using the CTAB method. The fingerprints of genotypes were screened with 58 simple sequence repeats (SSR) separated in the whole genome of *B. napus* (Supplementary Material S1). The SSR products was visualized on 10% polyacrylamide gel electrophoresis for genotyping, and the SSR bands were described by absence (0) or presence (1).

The DNA methylation was analyzed by methylation-sensitive amplification polymorphism (McClelland et al. 1994; Xiong et al. 2013). The DNA was digested by the combination of restriction enzyme *Msp* I + *EcoR* I and *Hpa* II + *EcoR* I, and then linked with adaptors for pre-amplified. The pre-amplification products were diluted tenfold and amplified by 14 selective-primers (Supplementary material S2). The products of selective amplification were visualized on 6.5% polyacrylamide sequencing gels on a Licor-4300 DNA analyzer.

## Acknowledgements

This study was partly supported by the China Postdoctoral Science Foundation (2016M582500), the Fundamental Research Funds for the Central Universities in China (XDJK2016C080), the Chongqing Postdoctoral Science Foundation (Xm2016029), the Science and Technology Innovation Project of Chongqing (cstc2015shms-ztzx80005, cstc2015shms-ztzx80007 and cstc2015 shms-ztzx80009)

**Supplementary material 1** SSR primers for genotyping of hexaploid. Distribution of loci amplified with simple sequence repeat primers. The primers were prefixed with BRAS/CB from Celera AgGen Brassica Consortium; with SWUC, PUT, CEN from Southwest University; prefix Na, Ra, Ol from BBSRC microsatelite programme (Lowe et al. 2003); with NIAB from The National Institute of Agricultural Biotechnology; with PUT from <u>http://www.plantgdb.org</u>; with sR, sN, sS from Agriculture and Agri-Food Canada; with BRMS from Suwabe et al. (2002); with CNU from Chungnam National University; with FITO from http://www.asbornlab.agronomy.wisc.edu/research/maps/ssrs.html; with BN were from Szewc-McFadden et al. (1996); with MR from Uzanova and Ecke (1999); with sOR from Qiu et al.(2006); with ENA from Choi et al.(2007); with CN from Long et al.(2007)

**Supplementary material 2** Sequence of primers for methylation sensitive amplified polymorphism

